# Human scalp hair as a thermoregulatory adaptation

**DOI:** 10.1101/2023.01.21.524663

**Authors:** Tina Lasisi, James W Smallcombe, W. Larry Kenney, Mark D. Shriver, Benjamin Zydney, Nina G. Jablonski, George Havenith

## Abstract

Humans are unique among mammals in having a functionally naked body with a hair-covered scalp. Scalp hair is exceptionally variable across populations within *Homo sapiens*. Neither the function of human scalp hair nor the consequences of variation in its morphology have been studied within an evolutionary framework. A thermoregulatory role for human scalp hair has been previously suggested. Here, we present experimental evidence on the potential evolutionary function of human scalp hair and variation in its morphology. Using a thermal manikin and human hair wigs at different wind speeds in a temperature and humidity-controlled environment, with and without simulated solar radiation, we collected data on the convective, radiative, and evaporative heat fluxes to and from the scalp in relation to properties of a range of hair morphologies, as well as a naked scalp. We find evidence for a significant reduction in solar radiation influx to the scalp in the presence of hair. Maximal evaporative heat loss potential from the scalp is reduced by the presence of hair, but the amount of sweat required on the scalp to balance the incoming solar heat (i.e. zero heat gain) is reduced in the presence of hair. Particularly, we find that hair that is more tightly curled offers increased protection against heat gain from solar radiation.

**Significance:** The evolution of human scalp hair might be explained by thermoregulation pressures experienced in hot and arid environments. Bipedal posture and a hairless body may have necessitated the development of scalp hair to minimize heat gain from solar radiation, particularly in hominins with large brains. We used a thermal manikin and human-hair wigs to examine this thermoregulatory hypothesis. We found that scalp hair reduces heat gain from solar radiation; tightly curled hair is most protective. Specifically, our results show that hair protects the scalp from solar radiation while minimizing the amount of sweat required to offset heat gain, with tightly curled hair providing the most protection.

## Introduction

Within evolutionary anthropology, there is particular interest in understanding traits that distinguish the human lineage from its closest living relatives. By studying the function of traits that are considered distinctly human, we can gain insight into evolutionary selective pressures that shaped the species.

Bipedalism, encephalization, and loss of body hair are three traits of prime interest in the study of hominin evolution and the emergence of the genus *Homo* (1, 2). Research on the evolution of these quintessentially human traits has attempted to identify the selective pressures that have shaped them. In each case, thermoregulation has been implicated as a potential contributing factor (1, 3).

Thermoregulation is important for all living organisms, but the human lineage evolved several physical and behavioral traits that may have posed new challenges to its basic primate physiological mechanisms of temperature regulation. Principally, the emergence of prolonged bipedal striding and running occurred at the same time (∼ 2 million years ago) as the evolution of larger brain size (4, 5). Thus, the costs of overheating due to the metabolic heat production associated with locomotion were multiplied by the increased heat sensitivity of a large brain. These new thermoregulatory challenges required new solutions.

Sweating works in tandem with a seemingly hairless body to create a highly effective cooling system. But this physiological response is not without cost. Sweating increases the need for fluid replacement and thus safe water, and can lead to dehydration, which poses a significant risk in modern conditions as well(6). Therefore, less costly adaptations for keeping cool (i.e., scalp hair) may have offered significant additional benefits to an encephalized hominin.

While the insulating function of hair in cold environments is intuitively appreciated, mammalian coats can also reduce total heat gain from the environment in hot conditions (7, 8). Many terrestrial mammals in hot climates have evolved (active) cooling mechanisms (e.g., respiratory cooling via the carotid rete) that do not require a hairless body. In doing so, these mammals were able to maintain both the passive thermoprotective benefits of hair as well as its other functions in physically protecting the skin.

Few evolutionary anthropologists have studied human scalp hair because, unlike other traits like bipedalism or encephalization, the function of scalp hair is less clear. Some scholars have speculated about the evolutionary pressures that may have led to the emergence (or retention) of scalp hair. The most testable hypothesis among these is that scalp hairs evolved for thermoregulation purposes. Yet, models investigating early hominins’ temperature regulation do not consider scalp hair explicitly.

In comparison, physiologists and environmental ergonomists have given some thought to the thermal effect of scalp hair. Specifically, a few studies have examined the impact of scalp hair on sweating and heat loss. Cabanac and Brinnel found that bald men sweat in that area at two to three times the rate of men with scalp hair(9).

Initially, this aligns with the logical idea that a hairless head would be better off in terms of heat loss because it has no barrier blocking evaporation. However, according to Coelho et al.’s more recent study from 2010(10), there may be a greater disadvantage to having no scalp hair since it also subjects the scalp to higher heat loads, mainly through solar radiation influx.

After conducting experiments involving ten men, Coelho and colleagues found that after men shaved their scalp hair, their head hair required more evaporative cooling during exercise in the sun. These findings challenge the widely accepted belief that a bald head is optimal for heat balance(3).

However, the scenario of hair/no hair is not always realistic. Shin et al.(11) found that people with shorter hair (5mm) lost heat quicker than those with longer hair 100-130mm). This helps explain the spectrum between a hairy and bald scalp. But one key question has yet to be answered: how does scalp hair morphology (or texture) affect thermal load?

Human populations exhibit a great deal of variability in scalp hair morphology(12–14). In addition, tightly curled hair—which is common in many African populations—is a uniquely human characteristic amongst mostly straight-haired non-domesticated mammals.

Jablonski and Chaplin(15) suggest that this distinctive phenotype may have an advantage in reducing heat gain from exposure to sunlight. Additionally, the ubiquity of tightly curled hair in a continent with unmatched genetic diversity suggests the role of scalp hair morphology deserves further attention.

Given the orthograde bipedal posture of hominins, the emergence (or retention) of scalp hair may have struck an optimal balance between maximizing heat loss across the large surface area of the body and minimizing solar heat gain on the small surface area of the scalp (1), directly over the brain. Tightly curled hair may provide an additional reduction in heat influx beyond the capacity of typically straight mammalian hair (15). Thus, the evolution of (tightly curled) scalp hair may represent part of an integrated evolutionary response to new thermoregulatory challenges faced by increasingly encephalized hominins.

In this paper, we examined the effect of human scalp hair on scalp heat gain from solar radiation and further explored the effect of hair morphology (specifically hair curl). We used a thermal manikin to generate empirical data on heat losses and heat gains of a scalp—nude, and with human hair wigs of various textures. Thermal manikins are designed to model heat transfer between human skin and the environment and are often used to test heat exchange through textiles and clothing configurations across a range of ambient temperatures, humidities, wind speeds and radiation levels (16). This biophysical approach allowed for the collection of heat transfer data on the thermal properties of hair without the noise that is typically introduced when measuring on human participants through their variable physiological responses under heat stress. Finally, we interpreted the results in light of the hypothesis that (tightly curled) scalp hair may have evolved as a result of selection on reduced heat influx on an increasingly large human brain.

## Results

We carried out a series of manikin experiments in a climate-controlled chamber to ascertain the dry and evaporative (wet) heat fluxes to and from a scalp with no head covering (nude), straight hair, moderately curled hair, and tightly curled hair (see experimental design and conditions in Figure 1). Specifically, we sought to understand the effect of hair (presence/absence and morphology), solar radiation, and wind speed on heat loss from the scalp. Subsequently, we present the results for a climate of ° (see supplementary information, SI Appendix 1, for detailed methods).

**Figure 1.**
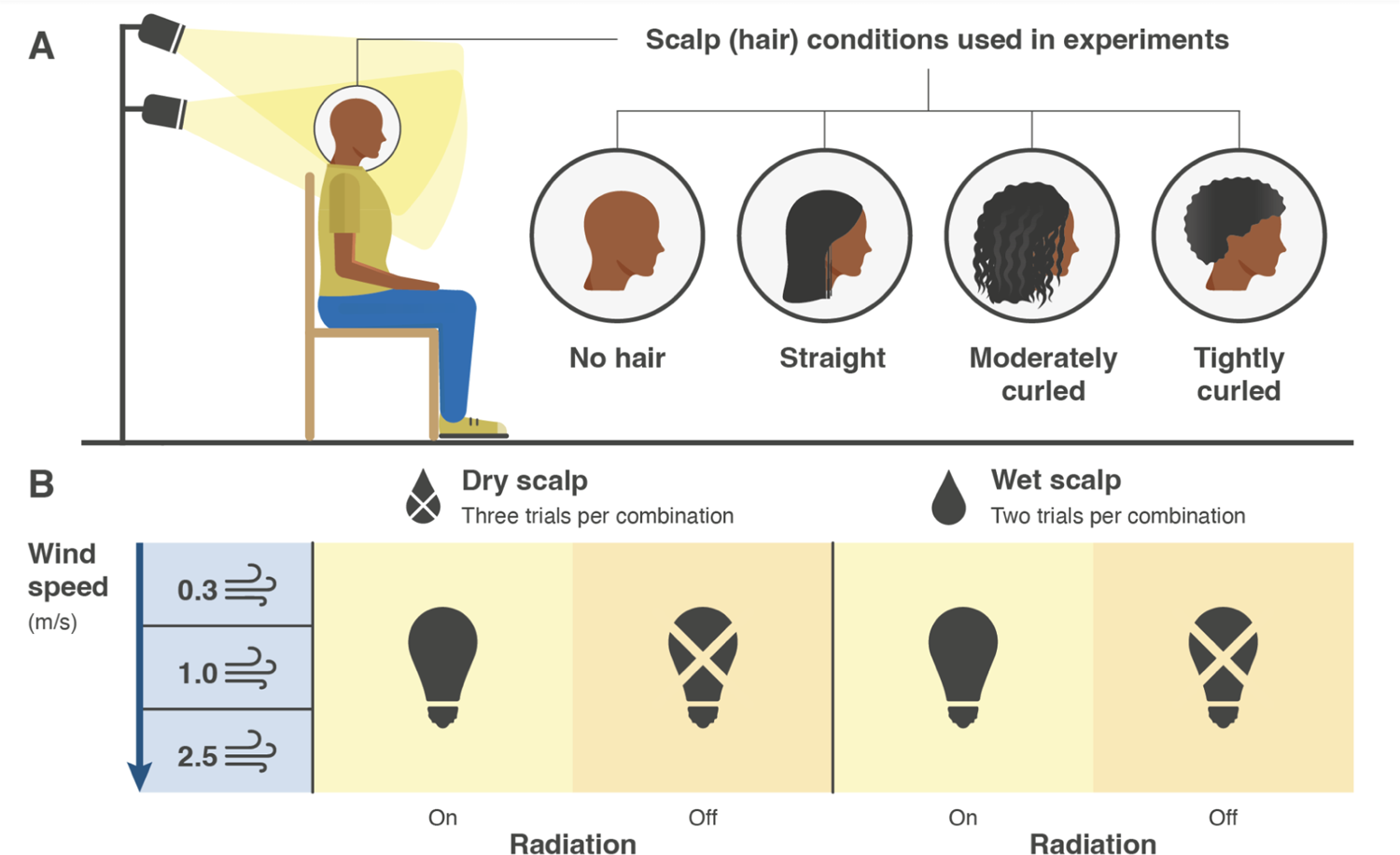
Diagram of the experimental setup and conditions. Panel A shows the physical setup and the four scalp-hair conditions simulated: none, straight, moderately curled, and tightly curled. Panel B shows the experimental input variables: dry or wet scalp, three wind speeds, and radiation (light) on or off.

For dry heat exchange under solar load, we found that in a climate with an ambient temperature of 30 °C, 60% rh, and solar radiation (∼788 W/m2), most conditions resulted in heat gain (negative heat loss), with the exception of the 1 m/s wind speed condition where tightly curled hair had some heat loss and at 2.5 m/s wind speed where both tightly and moderately curled hair showed some heat loss (see Fig 2). In general, the pattern we observed is that the highest solar heat gain was experienced under the nude condition and that straight hair, moderately curled hair, and tightly curled hair showed decreasing heat gain in that order.

**Figure 2.**
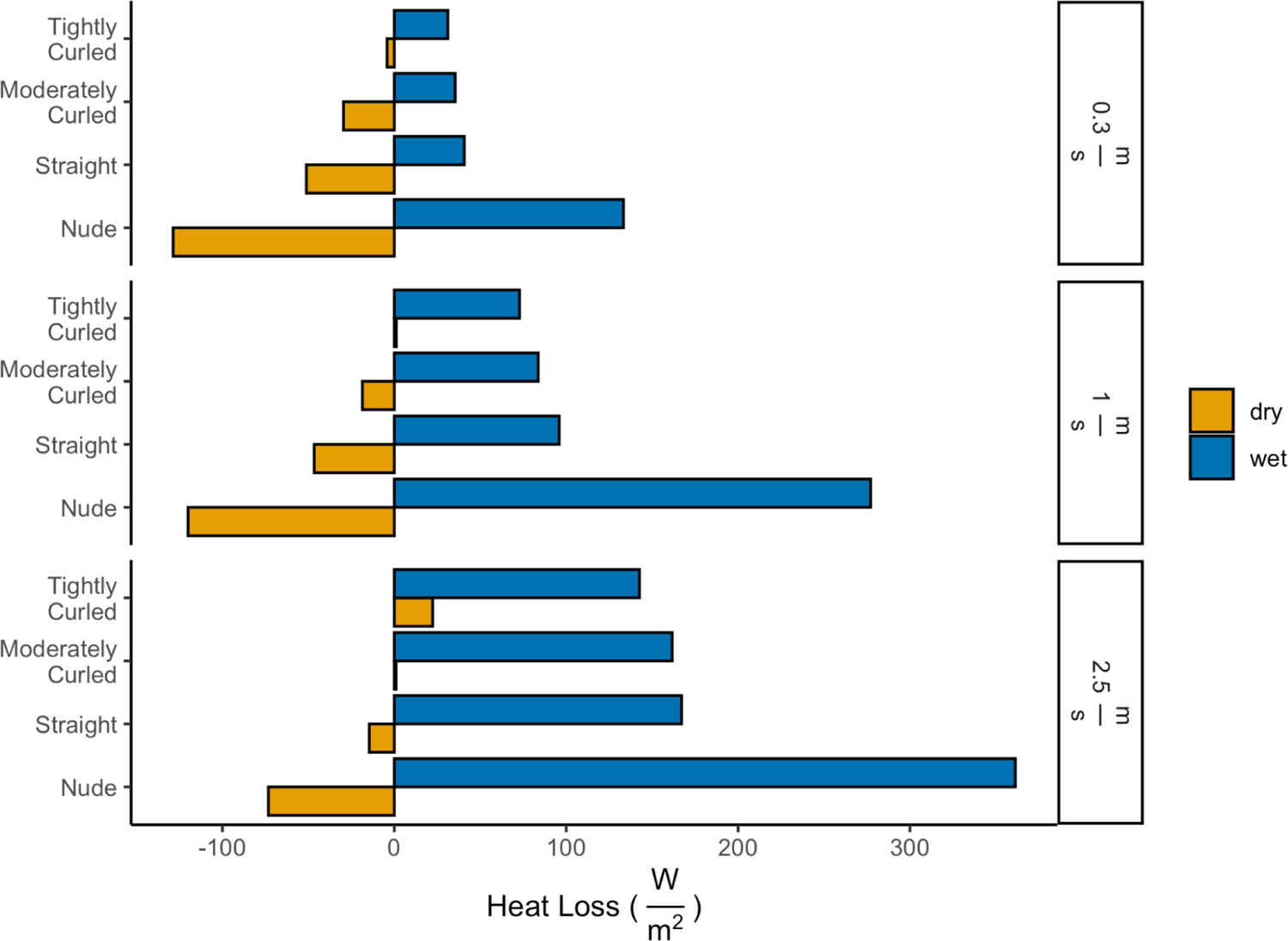
Net Dry heat exchange while exposed to a solar load with dry skin (yellow) and net wet/evaporative heat loss while exposed to solar load for a fully wet skin (blue) for all wig conditions and wind speeds were calculated for an ambient temperature of 30 °C with 60% relative humidity. Note: negative numbers indicate a heat gain; positive a heat loss.

For wet heat loss under solar load (see Fig 2), we see that the nude condition allows for much higher evaporative heat loss than any of the hair coverings. There is a small difference among the different wigs, with tightly curled hair decreasing wet heat loss more than other wigs. Overall, these results confirm that any type of barrier reduces the efficiency of evaporative heat loss and that evaporation is improved by improved convection associated with higher wind speeds.

We found the effect of wind speed to be a significant predictor of heat loss in both dry and wet skin conditions. The effect of different wigs was also a significant predictor of heat loss in the dry skin experiments while differences between wigs were not significant for wet skin heat loss (see supplementary results in SI Appendix 2).

Looking specifically at the effect of solar influx between our dry heat loss experiments and our evaporative heat loss experiments, we saw that solar heat gain was primarily a factor in dry heat loss conditions (see Fig 3a). In particular, we saw a large difference between the nude condition and any hair coverings, but there was also a clear pattern of decreased solar heat gain with increased hair curl. In our evaporative heat loss experiments, heat gain from solar radiation was highest in the nude condition at a windspeed of 0.3 m/s (see Fig 3b). However, there was no apparent distinction among wigs at that windspeed nor any difference between any hair coverings at other wind speeds.

**Figure 3.**
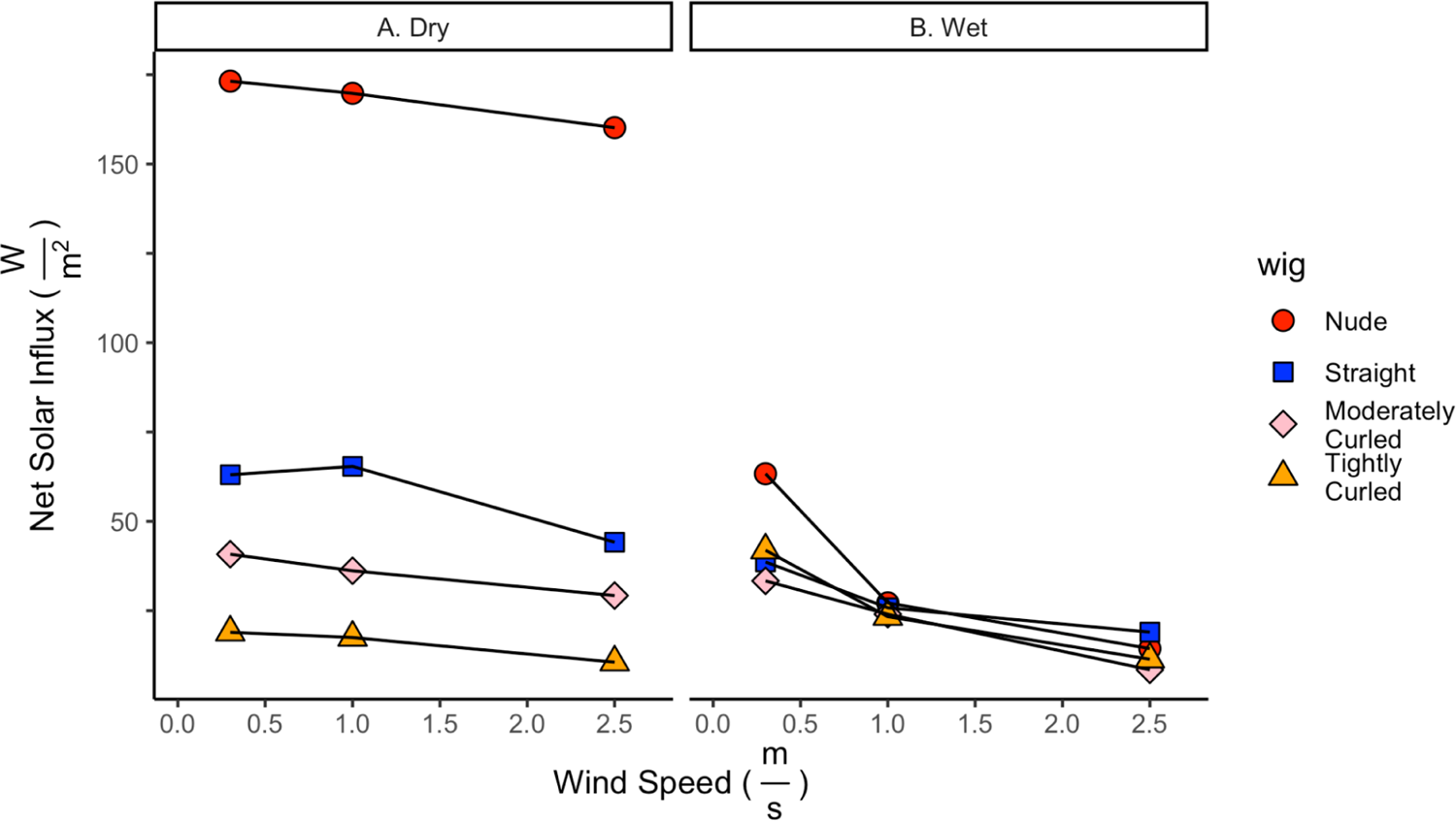
Net solar influx in the dry skin (A; radiant and convective heat exchanges) and wet skin (B; radiant, convective and evaporative heat exchanges)) condition in a *30 °C, 60% relative humidity condition*.

Based on our dry and wet data, we calculate the amount of sweat evaporation (maximum evaporative cooling potential) and the amount of sweat that would be required to achieve heat balance (zero heat gain for the scalp), under solar radiation (∼788 W/m^2^) at 30°*C*. We find that maximum evaporative cooling potential is highest with a nude scalp, followed by straight hair, moderately curled hair, and tightly curled hair. There is also an increase in evaporative cooling with increased wind speed (see Figure 4a). However, if we change from the ‘maximal heat losses’ paradigm to a paradigm where all heat influx from solar is exactly balanced by convective and evaporative heat losses from the scalp (heat balance) considering the net heat gain in each condition, tightly curled hair needed the least, and only minimal sweat to achieve heat balance at 0.3 m/s and none at higher wind speeds (see Figure 4b). Moderately curled hair and straight hair followed with an increased sweat requirement each. The nude scalp needed the most sweat to achieve heat balance (see Figure 4b). Wind speed reduced the amount of sweat required across head coverings due to the increased convective losses. See supplementary methods (SI Appendix I).

**Figure 4.**
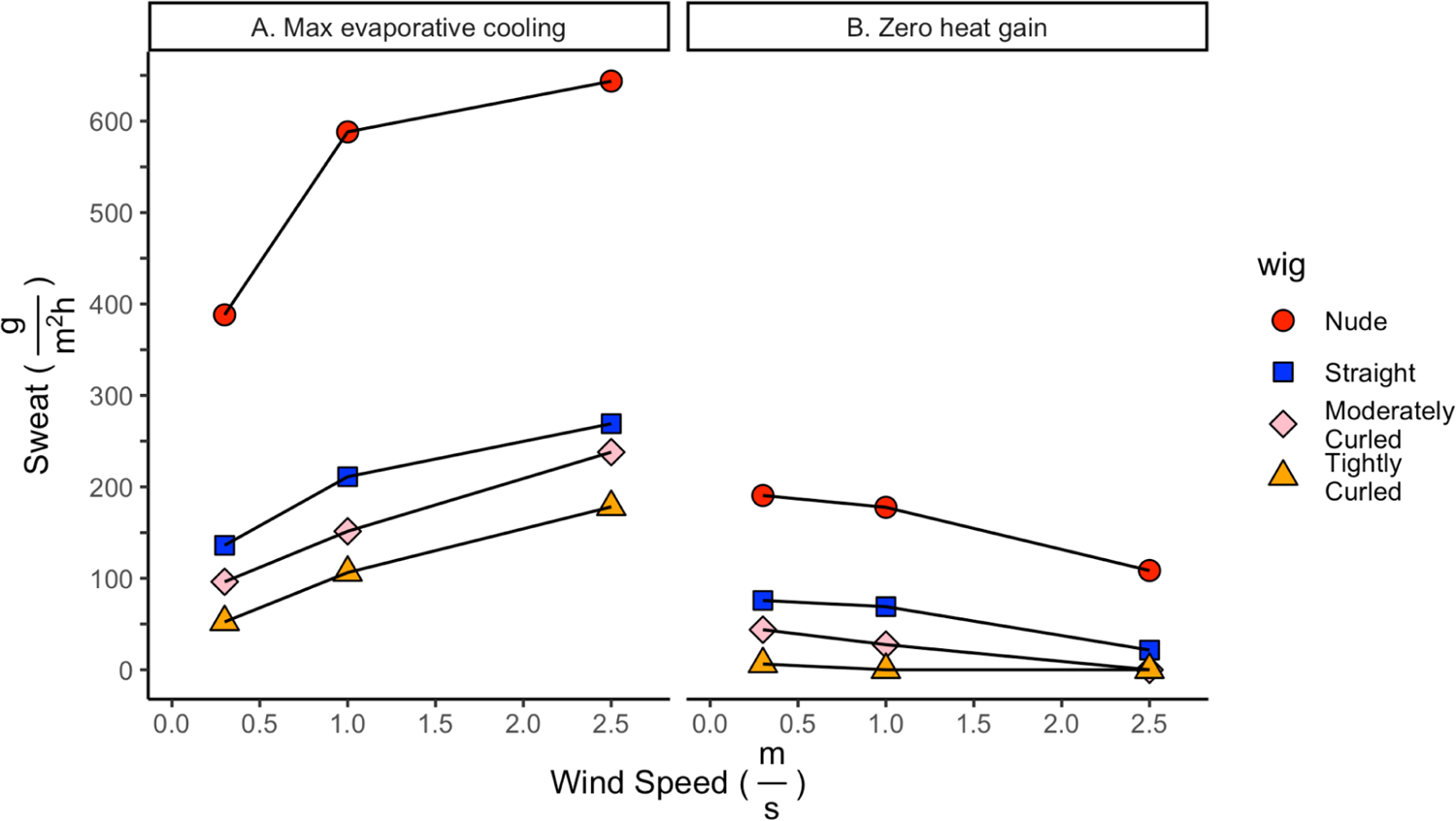
Estimated required sweat rate to achieve maximum evaporative heat loss potential from fully wet scalp skin in the climate (A) and sweat rate required to achieve zero heat gain, i.e., heat balance, (B) across different wig conditions and wind speeds at an ambient temperature of 30 °C and relative humidity of 60%.

## Discussion

Human scalp hair morphology exhibits considerable variation among human populations and includes phenotypes like tightly curled hair—a trait not found in any other non-domesticated mammal. Yet, we do not understand the evolutionary function of this trait. In this study, we employed an experimental approach using a thermal manikin and human hair wigs to examine a thermoregulatory hypothesis of scalp hair (morphology). Our results suggest that scalp hair reduces heat gain from solar radiation and that tightly curled hair, in particular, offers the most protection.

### Experimental evidence for thermoprotective function of human scalp hair

We measured dry and evaporative (wet) heat fluxes to and from the scalp to generate empirical data on the thermal properties of a nude scalp and ones covered in straight hair, moderately curled hair, or tightly curled hair, respectively, when exposed to solar radiation. Our findings confirm that, regardless of texture, hair acts as a barrier that decreases heat loss from the body (in this case, the scalp) to its surroundings. However, we find that in the context of solar radiation, hair functionally minimizes heat gain by reducing the amount of thermal radiation that reaches the skin surface. This is in line with studies of animal coats in areas of high solar radiation (17).

Our most striking observation is the clear pattern of decreased solar heat gain with increased hair curl (Figure 3a). The closest parallel that can be found in the animal literature is the decrease in solar heat gain with increased depth of a fur coat (8). Tightly curled human hair form does not lay flat on the scalp and therefore increases the distance between the surface of the hair and the surface of the scalp.

To better interpret the significance of these results, we used the dry and wet heat flux data (Figure 2) to calculate the maximum amount of “sweat” that would be evaporated from a wet, nude scalp as well as one fully wet, but covered by different hair textures for *T_ambient_* = 30*°C* and 60% relative humidity (chosen to represent a plausible climate scenario for early hominins). Our findings, unsurprisingly, show that a nude scalp can evaporate far more sweat than one covered in hair (Figure 4a), but at the cost of increased water loss. This is in line with results from human trials, which agree that scalp hair is associated with decreased sweating (9–11). Interestingly, we see a clear trend in decreased evaporative potential as hair curl increases, which has not been previously demonstrated.

Finally, we used the data to calculate how much evaporative cooling would be necessary for the different head covering conditions to achieve heat balance (i.e., zero heat gain or loss; sweat evaporation compensating for the solar influx; Figure 4b). In other words, how much sweat would one need to secrete and evaporate to prevent warming of the scalp? Here, it becomes evident that the higher evaporative potential of a nude scalp comes at a cost. Even though a scalp covered in tightly curled hair can evaporate less sweat, required evaporation is also less due to a lower solar heat influx. Moderately curled hair and straight hair follow in that order. Taken together, the dry and wet data indicate that tightly curled hair ultimately confers thermal advantage by minimizing overall heat influx when exposed to solar radiation.

### Hair and human thermoregulation in the literature

The general consensus in the literature is that scalp hair hinders evaporative cooling (3, 9–11), which is also supported by our results. Adding context to these findings, our study takes into consideration the effect of solar radiation and shows that the relative disadvantage in evaporative cooling potential is dwarfed by the reduction in solar heat gain associated with hair.

In contrast to previous studies, we do not focus on exercise-induced hyperthermia at various ambient temperatures. The results of our study offer a complementary view of the possible effect of solar radiation-induced hyperthermia.

To fully understand the role of human scalp hair in an evolutionary context, it would be important to synthesize information acquired from both these scenarios to paint a realistic picture of the thermoregulatory demands that may have been experienced by hominins.

In their 2010 study, Coelho and colleagues present an evaluation of sweat rate and body temperature in an outdoor setting under the sun(10). Although this method produces more uncontrolled variation than ours—due to variable physiological responses of human participants under heat stress—it better reflects the natural environment that we want to test from an evolutionary standpoint.

Coelho et al. demonstrated that although those with hair on their scalp experienced less sweat production, all other temperature measurements were not significantly different between the two conditions. One interpretation of those results is that those who sweat more because they have no hair on their heads require more evaporative cooling to keep their bodies cool in environments with high solar radiation. This is in line with our results.

The conditions under which humans evolved were such that solar radiation was high and free sources of drinking water were scarce. In these circumstances, evolution would have favored adaptations for water conservation. A plausible scenario could be the evolution of tightly curled hair that insulated against heat and reduced water loss while also extending how long individuals could engage in strenuous physical activity before needing a drink of fresh water.

### Limitations and future directions

Some limitations to the present study must be noted. First, this manikin study measures the static thermal properties of hair as a material but does not take into account physiological responses to heat stress, like changing sweat rates and changing skin temperatures. Moreover, we simulated “sweat” by saturating the manikin’s (cotton) skin with water, simulating the ideal case of fully wet skin and providing the maximal cooling potential data of fig. 2 and 4. This will require a high sweat rate (Fig 4a). In the context of radiation, sweat may evaporate more easily with the radiant load, and sweat rates may be insufficient to keep the skin fully wet, implying that the ‘heat balance’ condition of fig 4b is more likely. This may have consequences for the interpretation of evaporative potential.

Second, our wigs represented a narrow range of human hair variation due to cost and limited availability associated with the human hair market. Future studies should be designed to broaden the range of hair tested to account for a larger range of hair curl, as well as hair density, other aspects of morphology, and color. Lastly, we used a single intensity of solar radiation. Our findings would be strengthened by ascertaining the relationship between the intensity of radiation and hair morphology.

These results are important for researchers trying to understand the evolution of early hominins and later human populations, since they provide insight into the specific contexts where hair, particularly tightly curled hair, may have been advantageous. Tightly curled hair has often been inaccurately linked with “wool,” but our findings show that contrary to what this comparison might imply, curly hair does not trap heat close to the body like an insulation layer(18, 19). Instead, its advantage is in protecting against excessive heat exposure from solar radiation while still allowing sufficient heat loss.

Though we do not yet understand the extent to which scalp hair helps regulate whole-body temperature, this paper provides some valuable preliminary findings. This research represents a first step in understanding the connection between human scalp hair and thermal load to the brain and body.

### Evolutionary context for a thermoregulatory hypothesis of human scalp hair evolution

We do not yet know when exactly any scalp or hair-related adaptations may have occurred; however, it is still helpful to consider some potential scenarios. Specifically, we should question the order in which relevant traits appeared. For example, did hominins’ first development of miniaturized hair follicles lead to loss of their scalp hair along with the rest of their body hair? Was the appearance of scalp hair a later adaptation? Or was scalp hair retained during early *Homo* evolution and the loss of terminal body hair?

We might also question whether tightly curled hair and scalp hair evolved together or if one developed before the other. Furthermore, it would be essential to investigate whether this morphology only appeared once in humans or repeatedly through convergent evolution.

Based on our current knowledge of the hominin evolutionary history, a crucial period of time we may want to focus on is 2 million years ago, with the dispersal of *Homo erectus*, and 80 thousand years ago, with the dispersal of *Homo sapiens*. Between these two periods of time, we also suspect numerous dispersals leading to Neanderthal and Denisovan populations in Eurasia. Thus, an interesting point of consideration is whether the evolution of (tightly curled) scalp hair would have occurred prior to any hominin dispersals or prior only to *Homo sapiens’* dispersal.

The timing of scalp hair evolution should be considered in the face of two main factors: the evolution of “hairless” skin in response to thermoregulatory pressures and the evolution of larger brains that represented a thermoregulatory constraint. Our experimental results provide compelling evidence for a scenario whereby tightly curled scalp hair may have alleviated a thermoregulatory constraint on increased brain size (and increased thermoregulatory stress) with the passive heat load reduction offered by such hair.

Modeling the various anatomies of different hominins would be necessary to answer questions about timing or relative fitness advantage of (tightly curled) scalp hair. This modeling would need to take into consideration skeletal robusticity (which may have offered some benefits against heat gain, especially in the skull) and brain size (which might point to an increased risk of heat stroke).

Furthermore, how *Homo erectus*, Neanderthals, and *Homo sapiens* behaved would have been instrumental in their ability to change and adapt to different climates. If we take into account the relative importance of such behavioral adaptations, it would provide us with a better understanding of when scalp hair (curled or not) may have become an evolutionary advantage.

To our knowledge, this is the first report of the effect of human scalp hair morphology on solar heat gain. These results provide a good starting point for discussion and further research of the evolutionary function of human scalp hair (and variation in its morphology). The most pressing question to address is the significance of these findings in the context of a full-body thermoregulatory response. The head represents a small fraction of the body’s total surface area. We must investigate whether hair offers enough protection to significantly affect body-wide thermoregulation or whether the local protection it offers to direct brain heating is sufficient to constitute an adaptive benefit.

## Materials and Methods

Experiments were conducted in a wind tunnel in a climate-controlled chamber using a full-body thermal manikin (model “Newton,” Thermetrics, Seattle, Washington, USA). Data presented in this paper are only from the head zone of the manikin (out of 34 independently controllable zones on the manikin). Dry manikin measurements measured convective heat loss and solar influx. Wet manikin measurements were performed with the manikin skin and air temperature set to be identical, which means no convective heat loss is present. Only solar influx and evaporative heat losses are present in that case. By measuring manikin heat losses dry and wet and with and without radiation, convective, evaporative, and solar heat exchanges could be separated (20–23).

The wigs used in the experiment ranged from straight to tightly curled (see SI Appendix) and were all black human hair of Chinese origin made with 8” hair fibers. Solar radiation was simulated with two lamps reaching a net radiation of ∼788 W/m2. This was chosen to represent the solar load that might reasonably be experienced at ground level during midday sun. The obtained data on convective, radiative, and evaporative heat exchanges of the manikin in the different wig configurations were used to calculate the heat fluxes to and from the scalp for a climate of 30 °C, 60% rh.

See SI Appendix for detailed methods.

## Supporting information

SI_Extended materials and methods

SI: Extended results

## Data Availability

Manikin data are available on GitHub, https://github.com/tinalasisi/HairManikin2023.

## Funding

This research is supported by The Wenner-Gren Foundation (Gr. 9911) and the National Science Foundation (No. 1847845).

## Notes

https://github.com/tinalasisi/HairManikin2023

https://tinalasisi.github.io/HairManikin2023/analysis.html

### Competing Interest Statement

The authors have declared no competing interest.

### Summary of Updates

Some typographical and grammatical errors in the significance statement were fixed.

