## Supplementary material for "Human scalp hair as a thermoregulatory adaptation": SI Extended Methods

### SI: Extended materials and methods

#### Materials & Methods

##### Wigs

The global market of human hair limits feasible options for this project, so we decided to perform our experiments on three wigs made of untreated human hair reported to be of Chinese origin. A single wig is made of the hair of multiple individuals; however, in the process of production, hairs of uniform appearance are combined, resulting in less variation than would be observed across a natural scalp (Tarlo, 2017). To minimize variation between wigs, we used three naturally black human hair wigs of Chinese origin from the same source with 8” hair fibers (Figure 1).  The principle difference between these wigs is that one is straight, the second has artificially been made moderately curly, and the third has artificially been made tightly curled.

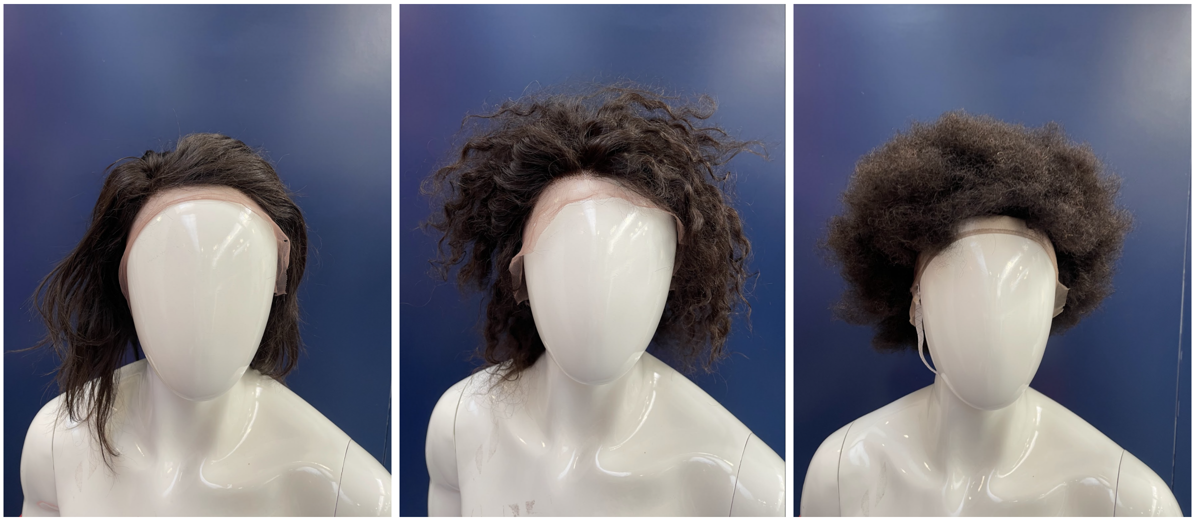

**Figure 1.** Wigs used in experiments. The wigs used in the experiment are made of human hair and were purchased from a purveyor who fashions different hairstyles out of “Chinese virgin hair”. The wigs are all made with hair fibers that are 8” long - the only difference is the tightness of the curl that was set in the wigs.

##### Equipment

The experiments used a thermal manikin (model “Newton,” Thermetrics, Seattle, Washington, USA). The “Newton” manikin is made of a shell of copper-filled carbon-epoxy and features embedded wire elements that heat the surface (skin) of the manikin (with a maximum heat output of 800 $W/m^2$). The manikin skin is typically heated to skin temperature around 34-35 °C (representative of a skin temperature during mild exercise in the heat).

In order to analyze radiant heat influx to the head as well as analyze convective and evaporative heat losses from the head, a thermal manikin was used in different combinations of solar radiation (788 W/m$^2$, on/off), different air temperatures, wind speeds, and either with wet (evaporative, convective and radiative heat exchange) or dry (convective and radiative heat exchange) skin(1–4). The manikin measures heat loss from its surface in 34 independently controlled body segments. For the present research question the focus was on the scalp.

In the dry skin experiments convective heat losses depend on the insulative properties of the wig and difference between the skin and air temperatures. When solar radiation is present the total heat loss from the manikin (convective loss minus solar influx) is reduced; for high solar fluxes heat loss may become zero or even negative, which would warm up the manikin and take it out of its control zone. So for the analysis/measurement of heat fluxes in the dry condition the air temperature is reduced to ensure a net heat loss of at least 20W/m$^2$ (as required by ISO 9920, (5)). The calculations take the different air temperatures into account (1–4). Measuring heat losses with and without solar radiation in the dry condition allows the calculation of the convective heat losses and, most importantly, the net solar radiation influx to the head.

For the wet skin testing air temperature is set equal to the manikin skin temperature. This stops any convective heat loss and thus the measured heat losses represent evaporative heat loss only (no solar radiation condition) or evaporative heat loss minus solar radiation influx (solar radiation condition).

All measurements were conducted in a climate-controlled chamber (TISS performance chamber, UK), in a custom-built wind tunnel. Climate data were registered with temperature sensors (Pt100) and humidity sensors (Vaisala).

Solar radiation was produced artificially using compact source iodide lamps (CSI; Thorn Lighting, Durham, UK) (6). The lamps filter out ionizing radiation to negligible values and thereafter produce a similar spectral content to that of sunlight. An array of two vertically aligned 1000-W metal halide lamps were placed posterior to the manikin. The average intensity of radiation across the scalp was 788 W/m$^2$, as measured immediately before and after each experiment (CM11 Pyranometer; Kipp and Zonen, The Netherlands). This level of Solar radiation is typical for what is observed under a clear sky during the typically hottest hours of the day for Eastern Africa(7).

Experiments were carried out at wind speeds of ~0.3m/s, 1m/s, and 2.5m/s, roughly comparable to air movement when still, walking, and running, respectively. A diagram and image of the setup are shown in Figures 2 and 3 respectively.

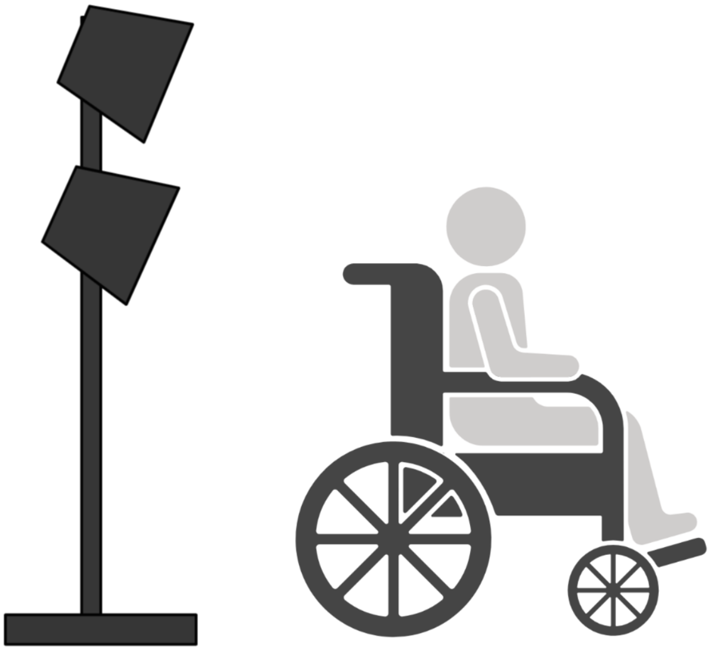

**Figure 2. Diagram of experimental setup.** The thermal manikin was sitting with its back toward the source of radiation.

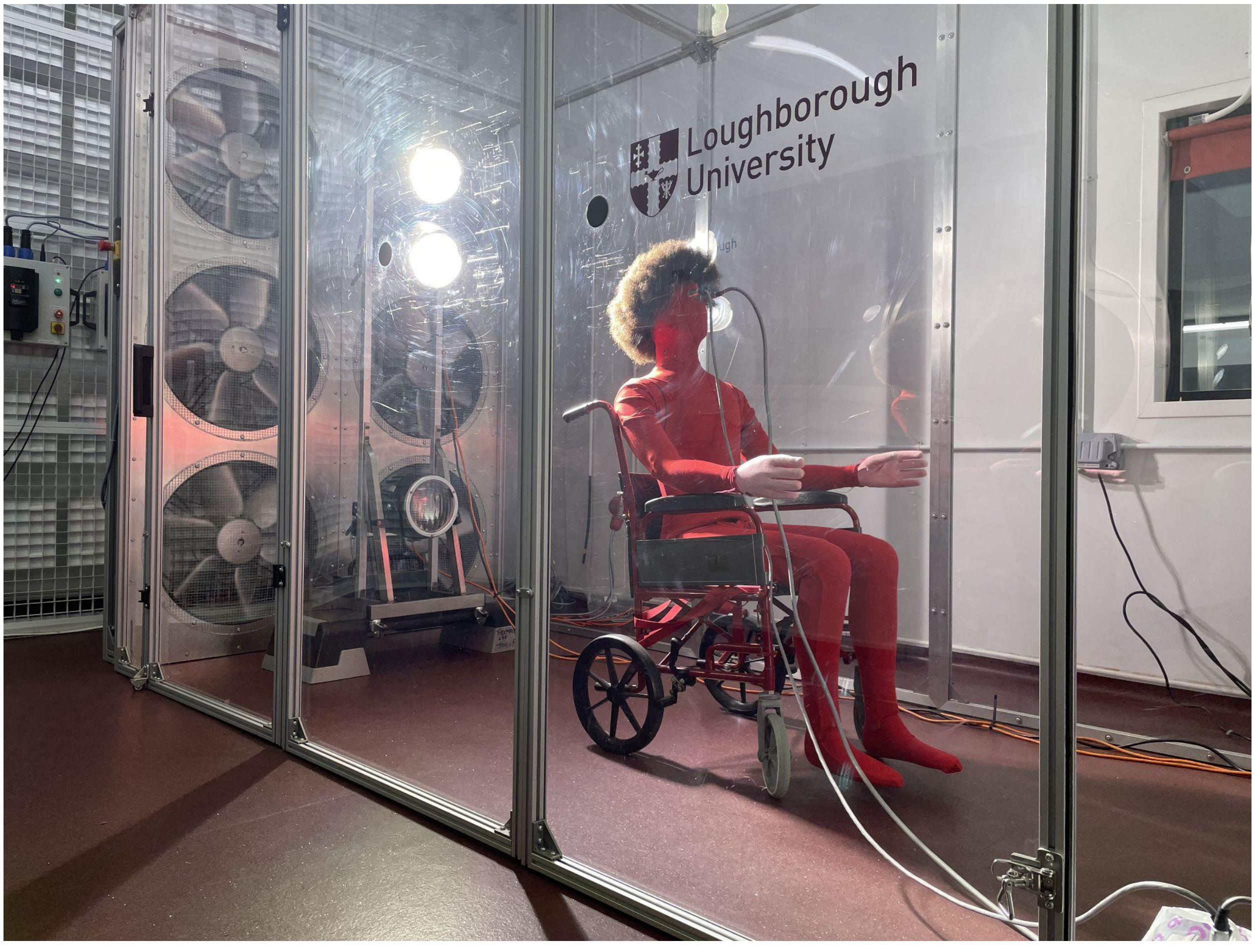


**Figure 3. Image of the experimental setup.** The manikin set-up inside the wind tunnel inside the climate-controlled chamber at Loughborough University.

The dry heat resistance experiments were carried out from October to November 2018, under the supervision of Dr. George Havenith and Dr. James Smallcombe at Loughborough University (Loughborough, UK). Air temperatures were set at 2 °C and 10 °C, depending on wind speed (ensuring the manikin lost at least 20 W/m2. The evaporative heat loss experiments were carried out from May to August 2019 by Dr. James Smallcombe at Loughborough University. The wet measurements involved completely saturating the manikin's cotton "scalp" and upper body with water from a spray bottle (simulating a heavy sweating scalp) and taking measurements with the room temperature settings $T_{manikin} = T_{ambient}$ to remove any convective heat losses. Humidity in the chamber during the wet testing was 48%.
To consider the outcomes for a relevant climate scenario to the geographical area for early *Homo* evolution, all heat exchanges were re-calculated for a $T_{ambient} = 30\,^{\circ}C$ and 60% relative humidity (rh) environment. This was associated with an expected skin temperature of 35 °C.

#### Analyses

##### Step 1: Processing manikin data

First, the raw manikin data was processed by Dr. George Havenith to convert the manikin readings to heat loss in W/m$^2$ for each region of the manikin; subsequently, the readings for each experiment over a period of stability were averaged for the head region.

The measurements in dry and wet skin conditions, both with and without solar simulation, allowed deduction of the individual heat fluxes (solar radiation influx; convective and evaporative outflux. Calculations of heat and vapor resistances of the nude head (surface air layer resistances) and the three wigs (wig + surface air layer resistances) subsequently allowed calculation of the individual heat fluxes at different air temperatures and humidities.

##### Step 2: Calculating net solar influx

To get the net solar influx, $I\ (\frac{W}{m^2})$, we calculated the difference in heat loss between the solar ($H_{Dry}^{Solar}$) and non-solar ($H_{Dry}$) experiments under the assumption that the net solar radiant influx remained constant (8).

$$I_{Dry} = H_{Dry} - H_{Dry}^{Solar} \qquad (\frac{W}{m^2}) \quad (3)$$

$$I_{wet} = H_{wet} - H_{wet}^{Solar} \qquad (\frac{W}{m^2}) \quad (4)$$

##### Step 2: Calculating thermal resistances

The dry heat resistance, $I_t\ (\frac{m^2K}{W})$, and evaporative heat resistance, $R_{e,t}\ (\frac{m^2kPa}{W})$, of the nude head (surface air layer insulation) and the wigs (including the outer surface air layer insulation) were then calculated in the non-solar condition as follows, with heat losses $H_{dry}$ and $H_{wet}$ in $\frac{W}{m^2}$:

$$I_t = \frac{T_{Skin} - T_{Air}}{H_{Dry}} \qquad (\frac{m^2K}{W}) \qquad (1)$$

$$R_{e,t} = \frac{P_{H2O,Skin} - P_{H2O,Air}}{H_{Evap}} \qquad (\frac{m^2kPa}{W}) \qquad (2)$$

where $P_{H2O,Skin}$ and $P_{H2O,Air}$ are the saturated water vapor pressure at the skin and the water vapor pressure in the environment during the testing, respectively, both measured in $kPa$.

#### Step 4: Adjusting for temperature and humidity

In order to evaluate the heat losses under solar exposure in a tropical environment, the heat losses were recalculated to 30°C, 60% rh using the obtained data assuming a skin temperature of 35°C:

$$H_{Dry}^{30^{\circ}C} = \frac{35 - 30}{I_t} \quad (\frac{W}{m^2}) \quad (5)$$

For dry skin with solar radiation, $H_{Dry}^{30^{\circ}C, Solar}(\frac{W}{m^2})$ is then calculated as the temperature adjusted convective heat loss minus the dry solar radiation influx (minus, since it is an influx rather than a loss):

$$H_{Dry}^{30^{\circ}C, Solar} = H_{Dry}^{30^{\circ}C} - I_{Dry} \quad (\frac{W}{m^2}) \quad (6)$$

For wet skin with solar radiation, $H_{Wet}^{30^{\circ}C, Solar}(\frac{W}{m^2})$ is the sum of the evaporative and dry heat losses minus the dry solar radiation influx,

where the evaporative heat loss is calculated as:

$$H_{evap} = \frac{5.63 - 2.53}{R_{e,t}} \quad (\frac{W}{m^2})(7)$$

Where 5.63 is the vapor pressure of wet skin at 35°C and 2.53 kPa is the vapor pressure in the ambient air at 30 °C, 60% rh. As this vapor pressure is the same as the one used in the wet data collection, those actual data can be used and a correction is not required. This then allowed calculation of the total heat losses at 30 °C, 60% rh in the sun, with wet skin:

$$H_{Wet}^{30^{\circ}C, Solar} = H_{Evap + Dry}^{30^{\circ}C, Solar} = H_{Evap} + H_{Dry}^{30^{\circ}C, Solar} - I_{Dry} \quad (\frac{W}{m^2}) \quad (8)$$

#### Step 5: Calculating evaporative potential

The evaporative potential, $H_{Max}$, is the maximum value of evaporative heat loss for the different wigs and the nude condition, possible under the chosen solar radiation of 788 $\frac{W}{m^2}$ assuming a sustained, fully wetted skin at an ambient temperature of 30 °C, 60% rh (8). This is the difference between the calculated adjusted wet and dry heat losses with solar radiation. Note that values for dry heat loss under solar conditions at 30 °C tend to be negative, as solar heat influx overpowers convective losses.

$$H_{Evap,\,Max}^{30^{\circ}C,\,Solar} = H_{Wet}^{30^{\circ}C,\,Solar} - H_{Dry}^{30^{\circ}C,\,Solar} \qquad (\frac{W}{m^2}) \quad (9)$$

#### Step 6: Calculating sweat requirements needed to achieve $H_{Max}$

The sweat rate that is required to achieve this maximal potential evaporative heat loss ($H_{Evap,\,Max}$), i.e. $Sweat_{Max}$ $(\frac{g}{m^2h})$, at 30 °C is calculated using the heat of vaporization of sweat ($2430\ \frac{J}{g}$) and the conversion factor of $3600$ $(\frac{s}{h})$ (9):

$$Sweat_{Max} = \frac{H_{Evap,\,Max}^{30^{\circ}C,\,Solar} \cdot 3600}{2430} \qquad (\frac{g}{m^2h}) (10)$$

Finally, the amount of sweat required to exactly balance out the incoming solar load (i.e. zero heat loss or gain of the head from the environment), $Sweat_{Zero}$ $(\frac{g}{m^2h})$, is calculated as follows, with the logical requirement that we can never have negative sweat.

$$IF\ H_{Dry}^{30^{\circ}C,\,Solar} < 0,\ Sweat_{Zero} = -\frac{H_{Dry}^{30^{\circ}C,\,Solar} \cdot 3600}{2430}$$

$$ELSE,\ Sweat_{Zero} = 0 \qquad (11)$$

Analyses with data can be found at https://tinalasisi.github.io/HairManikin2022/analysis.html


### Figure legends

Figure 1. Wigs used in experiments. The wigs used in the experiment are made of human hair and were purchased from a purveyor who fashions different hairstyles out of “Chinese virgin hair”. The wigs are all made with hair fibers that are 8” long - the only difference is the tightness of the curl that was set in the wigs.

Figure 2. Diagram of experimental setup. The thermal manikin was sitting with its back toward the source of radiation.

Figure 3. Image of the experimental setup. The manikin set-up inside the wind tunnel inside the climate-controlled chamber at Loughborough University.
